# Mass cytometry reveals immunological response to radiation-induced cardiac fibrosis in mice

**DOI:** 10.1101/2021.02.23.432001

**Authors:** Yi Tang, Bing Wang, Mingjiao Sun, Pan Liu, Xue Zhang, Mingliang You, Bing Xia

**Affiliations:** Department of Radiation Oncology, Key Laboratory of Clinical Cancer Pharmacology and Toxicology Research of Zhejiang Province, Affiliated Hangzhou Cancer Hospital, Zhejiang University School of Medicine, Hangzhou 310002, China; Hangzhou Cancer Institute, Key Laboratory of Clinical Cancer Pharmacology and Toxicology Research of Zhejiang Province, Affiliated Hangzhou Cancer Hospital, Zhejiang University School of Medicine, Hangzhou 310002, China; Department of Pathology and Pathophysiology, and Department of Respiratory Medicine at Sir Run Run Shaw Hospital, Zhejiang University School of Medicine, Hangzhou, 310058, China

**Keywords:** Radiation-induced cardiac fibrosis (RICF), mass cytometry, immune response, macrophage

## Abstract

**Background:** Radiation-induced cardiac injury results in complex clinical presentations, unique management issues, and increased morbidity and mortality. But the underlying mechanism of radiation-induced cardiac injury is still unclear. The aim of this study is to explain the compositional alterations of innate and adaptive cells after thoracic irradiation and the relationship of macrophage with other adaptive cells.

**Methods:** C57BL/6 mice were anesthetized and performed with 20Gy single dose cardiac irradiation. We carried out mice echocardiography to examine cardiac function. Masson and immunohistochemical staining were adopted to analyze cardiac fibrosis. Blood and cardiac tissue were collected 7 days, 4 weeks and 8 weeks after irradiation and sham-irradiated mice were set as a control group. Part of blood and tissue samples were analyzed with mass cytometry. We also used remaining blood samples to measure pro-fibrotic cytokines.

**Results:** We observed reduced cardiac diastolic function and pathologically confirmed cardiac fibrosis after thoracic irradiation. Through mass cytometry, we identified that the proportion of neutrophils, macrophages and monocytes elevated. The ratio of T and B cells decreased after irradiation in the blood samples. For tissue samples, the proportion of macrophages, monocytes and neutrophils increased, endothelial cells reduced, but T and B cells also increased. The level of TGF-β, TNF-α, IL-6 climbed considerably, all of them are closely associated with the evolution of macrophage function during recovering phase of cardiac injury. In the correlation plot, we also discovered that CD4+T and CD8+T cells were strongly positively correlated with macrophages.

**Conclusion:** We illustrated the immunological response after radiation-induced cardiac fibrosis, and macrophage may play a crucial role during the process. The results might have therapeutic potential for targeting macrophages in order to reduce radiation-induced cardiac fibrosis.

## Introduction

Radiotherapy is an important treatment modality for cancer, previous literatures concluded that 50-70% cancer patients require radiotherapy intervention during the course of treatment. However, radiation-induced cardiac fibrosis (RICF) becomes an increasingly common complication that may occur following thoracic radiotherapy. The damage can manifest either long time after irradiation leading to severe cardiovascular symptoms or acutely in the days or weeks after radiation treatment(1). After ionizing radiation, cardiac endothelial activation occurs, causing the cells to switch towards a pro-inflammatory phase. Meanwhile, platelets accumulate to inflamed endothelium, release tumor necrosis factor-β1 (TGF-β1) and result to the inhibition of systhesis of endothelial thrombomodulin(TM)(2). The deficiency in TM leads to aggregation of thrombin, a powerful procoagulant, causing deposition of fibrin in the extravascular space surrounding capillaries and in the subintimal space of arteries and veins. Besides, with the elevated TGF-β1, neutrophils, macrophages and monocytes accumulate, together with lymphocytes, release cytokines that stimulate fibroblasts migration and proliferation to myofibroblasts. Myofibroblasts produce collagens and other extracellular matrix (ECM) components, contributing to fibrosis(2, 3). During this cardiac fibrotic states, immunologic phenomenon is considered to exert an important impact on post-irradiation injury. Previous investigations reveal that monocytes and macrophages mediate the fibrotic response through the production of cytokines, chemokines and alteration of extracellular matrix(4). Besides, T cells contribute both directly and indirectly to ECM remodeling and fibrosis, the activity of T cells influence monocyte recruitment and macrophage activation(5). Infiltrating inflammatory cells also interact with fibroblasts or other cells, promote myofibroblast formation and modulate the activities of myofibroblasts(6). However, a comprehensive analysis of changes across all immune populations after cardiac irradiation has not yet been undertaken.

Time-of-fight mass cytometry(CyTOF) is a new approach for the detection of cell markers at single cell level. With CyTOF, antibodies are uniquely labeled with isotopically pure metals instead of fluorophores and then quantified by inductively coupled plasma mass spectrometry. Compared with traditional flow cytometry, mass spectrometer provides exquisite resolution between detection channels, high multiplicity of biomarker detection, absolute quantification, no sample matrix effects, simplified measurement protocols and overall lower sample and reagent consumption(7). CyTOF was used to explore diversified immune responses in multiple inflammatory diseases, e.g. rheumatic diseases(8), atherosclerosis(9) and HIV infection(10). But there have been few reports on the application of CyTOF technology in the field of radiation-induced injury. Here we performed a model of radiation-induced cardiac fibrosis in mice and utilized CyTOF to describe the compositional changes of immune and endothelial cells before and after cardiac irradiation.

## Materials and Methods

### Animals

Eight- to 12-week-old male C57BL/6 mice (SPF grade, inbred strains, 20-25 g in weight) were purchased from Zhejiang Chinese Medical University Laboratory Animal Research Center and randomly assigned to cardiac irradiated and control groups. Mice were housed in specific-pathogen-free conditions, the room temperature was maintained at 22-25°C and humidity level at 30-45%, with access to a standard diet and water ad libitum. All animal studies were performed in accordance with an animal use protocol approved by the Animal Care ethics board of Zhejiang Chinese Medical University.

### Small animal radiation research platform (SARRP) for cardiac irradiation (CIR)

In short, adult C57BL/6 mice were anesthetized with 1% sodium pentobarbital 75 mg/kg intraperitoneally and lied on their back on the apparatus. In order to establish a mouse model of exploring the radiation cardiac injury in an organ-specific manner, SARRP(XStrahl Medical and Life Sciences, USA) was applied to target murine heart and minimize radiation dose to the surrounding organs (lungs and spinal cord). Before cardiac radiation, on-board cone beam computed tomography (CBCT) was performed within the system with 50 KV X-ray at 0.8 mA and filtered with 1.0 mm aluminum(Figure 1). And a 10 mm circular collimator was adopted to locate the radiation portal to the heart. The radiation plan was designed as one rotating-beam angled from 120 degree to -120 degree clockwise to simulate partial single arc radiotherapy. Whole heart irradiation was implemented using isocentric ARC-field irradiation with 220 kv X-ray, 13 mA, SSD 345 mm and a 0.15 mm Cu filter. A single dose of 20 Gy or 0 Gy (sham-irradiation) was administered to the whole heart. The field size was 10 mm×10 mm. The blood and whole heart samples from radiation group were collected at 7 days, 4 and 8 weeks after cardiac irradiation, non-irradiated samples were also collected as controls. Considering our time of sample collection after cardiac irradiation (7 days, 4 weeks and 8 weeks), we used 3 experimental groups and 1 control group, with 12 mice in each group.

**Figure 1.**
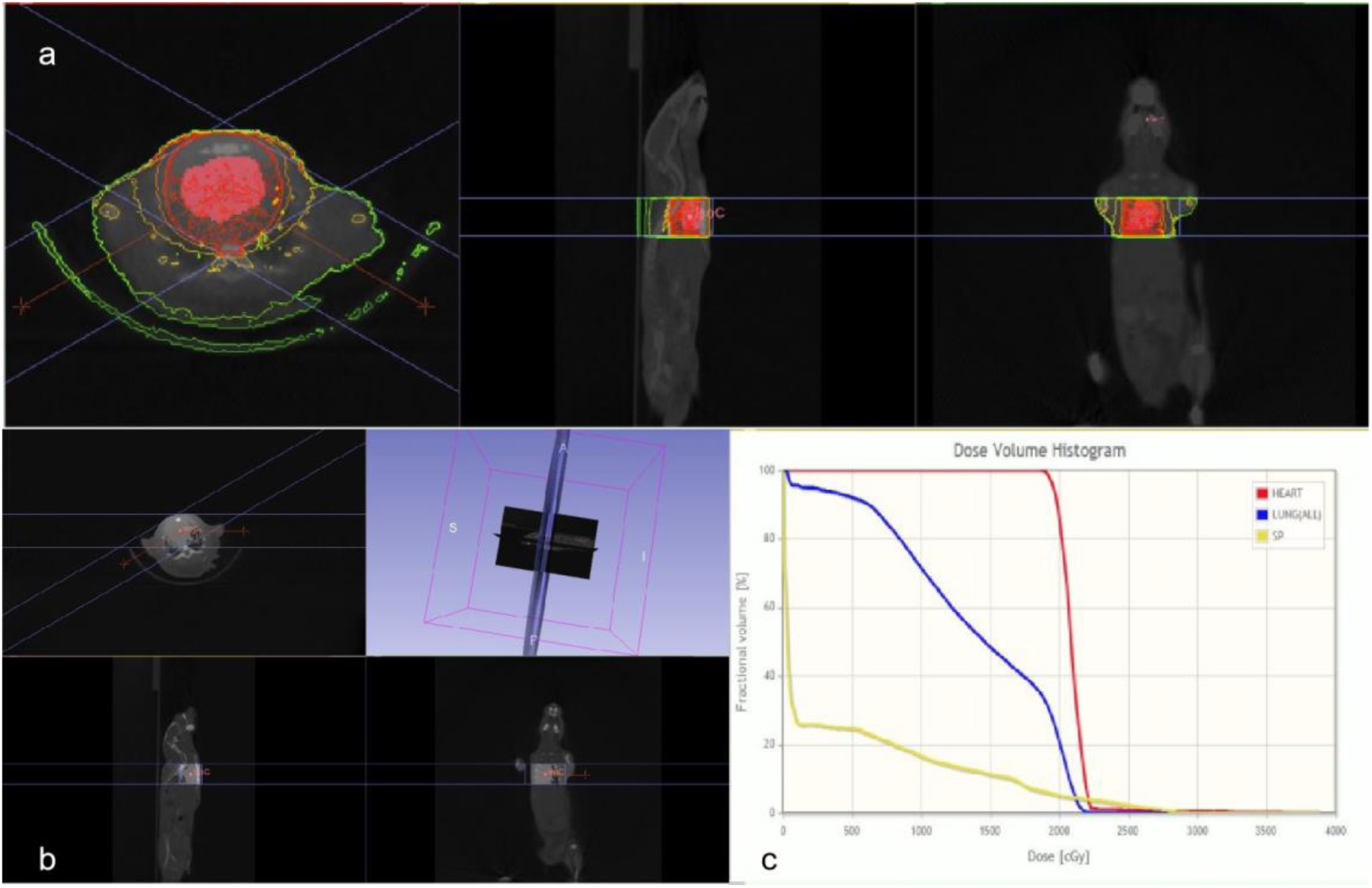
Establishment of a mouse model for studying the toxicities from cardiac irradiation.(a)shows transverse, sagittal and coronal plane during RIHD model. (b) Core beam computerized tomography before cardiac irradition. (c)shows representational Dose-Volume Histogram (DVH), the doses for lung and spinal cord were limited.

### Antibody labeling

A mass cytometry panel of 42 metal isotope-tagged antibodies (supplementary table1) was adopted to assess the immune and endothelial cells in the blood and tissue samples of mice. The antibodies were conjugated to the indicated metal tags using a MaxPAR X8 Antibody Conjugation Kit (Fluidigm, San Francisco, USA) according to the manufacturer’s protocol. The conjugated antibodies were diluted with Candor Antibody Stabilizer (Sigma) and titrated to the optimal concentrations.

For mice blood samples, we collected the blood from 9 mice in each timing for 3 samples to obtain sufficient cells for reliable mass cytometry. After lysis of the erythrocyte by ACK lysis buffer, samples were washed by fluorescence-activated cell sorting (FACS) buffer(PBS with 0.5% BSA and 0.02% NaN3) and kept at 4°C or on ice. For mice tissue samples, heart tissue was digested with DNAase, collagenase IV under 37°C for 1h. Then the digested fluid was washed with FACS buffer and centrifuged to leave the precipitation. After lysis of the erythrocyte by ACK lysis buffer, the precipitation was washed again with FACS buffer.

The pooled cells were then incubated with cisplatin for live or dead cell discrimination for 5 min. Following this, samples were blocked on ice for 20 min. Then the cells were stained with a mixture of metal-tagged antibodies (Supplementary table 1) targeting the surface antigens for 30 min. The cells were then washed twice with FACS buffer and incubated in Ir nucleic-acid intercalator (Fluidigm) in Fix and Permeabilization buffer (Fluidigm Sciences) at 4°C overnight. Then cells were stained with a mixture of metal isotope-conjugated intracellular antibody mix (Supplementary Table 1) for 30 min on ice. Following the washing of FACS buffer, the cells were counted and resuspended in distilled water filtered through capFACS tubes (Corning). Mass cytometry data were obtained using a Helios system (Helios, Fluidigm, South San Francisco, CA, USA). The raw data acquired was uploaded to a Cytobank web server (Cytobank Inc.) for further data processing and for gating out of dead cells and normalization beads.

Raw mass cytometry data were manually gated as live, singlet, and valid immune cells. The data were subjected to the metal isotope beads normalization method. The gated cell populations were clustered using the X-shift algorithm in MATLAB. The signal intensities of the markers were transformed using Arcsinh with a cofactor of 5. The percentage of each cluster was calculated as the percentage of all live cells. Clusters with cell frequency less than 0.1% have been ignored. To visualize the high-dimensional data in two dimensions, data from 10,000 randomly selected cells from each sample were processed using the nonlinear dimensionality reduction algorithm t-Distribution Stochastic Neighbor Embedding (t-SNE). Visualization of stochastic neighbor embedding (viSNE) was used to distinguish and gate major subgroups in peripheral blood of mice, which was generated with MATLAB (MATLAB Release 2015b macOS 64-bit version, MathWorks, Inc., Natick, MA, USA). viSNE is a clustering algorithm that transforms single cell high-dimensional cytometry data into a two dimensional visual representation. The heat map visualization was performed in Cytobank, the color gradient represents the various expression intensity between different types of markers.

### Histological Analysis

Murine hearts were collected for masson’s trichrome staining and immunochemistry analysis. Hearts were extracted and fixed in 10% formalin, and paraffin embedded. These paraffin-embedded samples were sectioned into 5um thick slices perpendicular to the apex to make sure both ventricles fully manifested. To detect the expression of vimentin, the paraffin-embedded heart sections were blocked in 3% H2O2, stained with primary antibody against vimentin (1 : 50) at 4°C overnight, and then incubated with peroxidase-coupled secondary antibody and DAB as a substrate. Images were captured at 400x magnification.

### Echocardiography

Irradiated and mice and non-irradiated control group were performed M-mode echocardiography 8 weeks after cardiac irradiation. Mice were first immobilized under isoflurane anesthesia, and acoustic coupling gel was applied to the chest after hair removal. We adjusted gains to eliminate background noise and 5 to 10 cycles were continuously recorded to ensure accurate readings.

### Specimen and Assay

Mice blood samples were collected with K2EDTA (dikalium salt of ethylenediaminetetraacetic acid) as the anticoagulant before and 4 weeks after cardiac irradiation. Blood samples were stored at 4°C immediately after collection, centrifuged within 6 hours of collection at 3000g for 30 min (4°C), and supernatants were collected and stored at-80°C before use. We tested 3 cytokines in the plasma samples. The serum levels of TGF-β, tumor necrosis factor-α(TNF-α), interleukin- 6 (IL-6) were determined using enzyme immunoassay (EIA) kits (R&D Systems, Minneapolis, MN, USA). The concentration of the above cytokines in culture medium was determined by comparing their optical density to the standard curve.

### Statistical Analysis

For the comparison of cell cluster frequencies, unpaired t-test was performed using SPSS 22.0(IBM Corporation, Armonk, NY, USA). P values <0.05 were considered statistically significant. All values are expressed as mean±SEM.

## Results

### Reduced cardiac diastolic function after irradiation

After cardiac irradiation, 8 weeks’ group had 16.7%(2/12) acute mortality compared with 0% in control and 4 weeks’ group. Autopsy of the two mice revealed no obvious cardiac abnormality. To determine whether cardiac irradiation result to impaired cardiac function, we performed echocardiography. Parameters of ejection fraction(EF), fraction shortening(FS), left ventricular volumn at end-systole(LVV-s), left ventricular volumn at end-diastole(LVV-d) were measured. As shown in Figure 2, mice after cardiac irradiation exhibited a significant decline in LVV-d (ul) compared with non-irradiated mice(LVV-d= 53.538±8.742 versus 83.192±8.962, p=0.015). We also found decreased value of EF (42.802%±18.304% versus 60.836%±8.918%, p=0.200) and FS (21.198%±10.922% versus 32.130%±6.545%, p=0.211), but they are not statistically significant. No difference was observed for LVV-s (ul) between irradiated and non-irradiated groups(22.643±12.454 versus 21.215±3.269, p=0.857).

**Figure 2.**
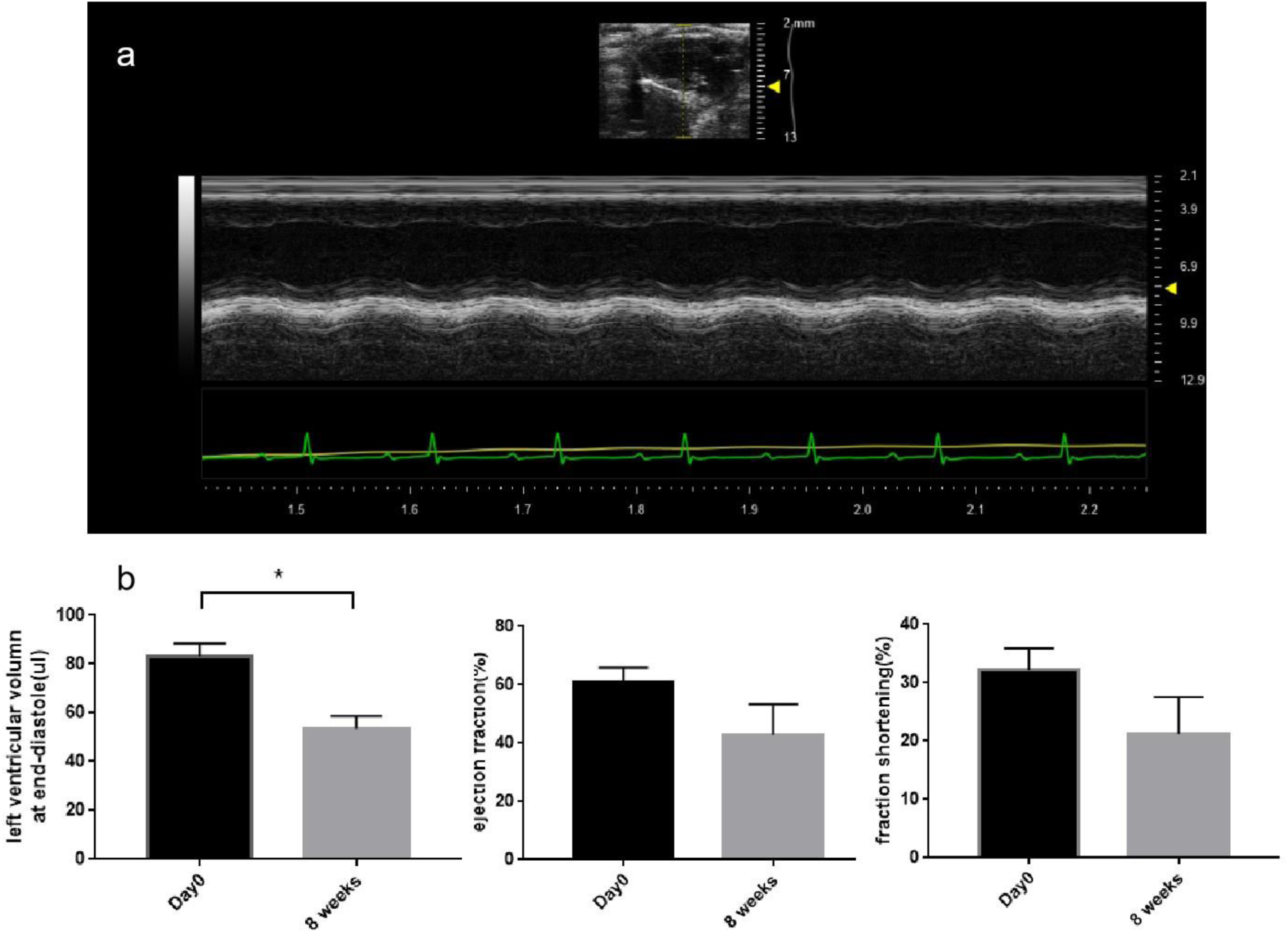
(a) M-mode echocardiography for mice after cardiac irradiation. (b) LVV-d declined 8 weeks after cardiac irradiation. The value of EF and FS decreased but the differences were not statistically significant. Columns represent the mean value of each group, bars±SEM, *p < 0.05.

### The model of radiation-induced cardiac injury

With the intent to reveal the pathological changes after cardiac irradiation, we selected murine cardiac tissue samples 8 weeks after irradiation for immunohistochemical analysis and masson’s trichrome staining. Non-irradiated heart was used as a negative control. It is clearly observed in masson’s trichrome staining that irradiated tissue had more collagen deposits in the myocardial interstitium compared with control group. Vimentin, which is a protein of intermediate filament family, labels fibroblasts with great sensitivity(11). Immunostaining revealed that vimentin was minimally expressed in non-irradiated cardiac samples, but was clearly upregulated 8 weeks after cardiac irradiation(Figure 3).

**Figure 3.**
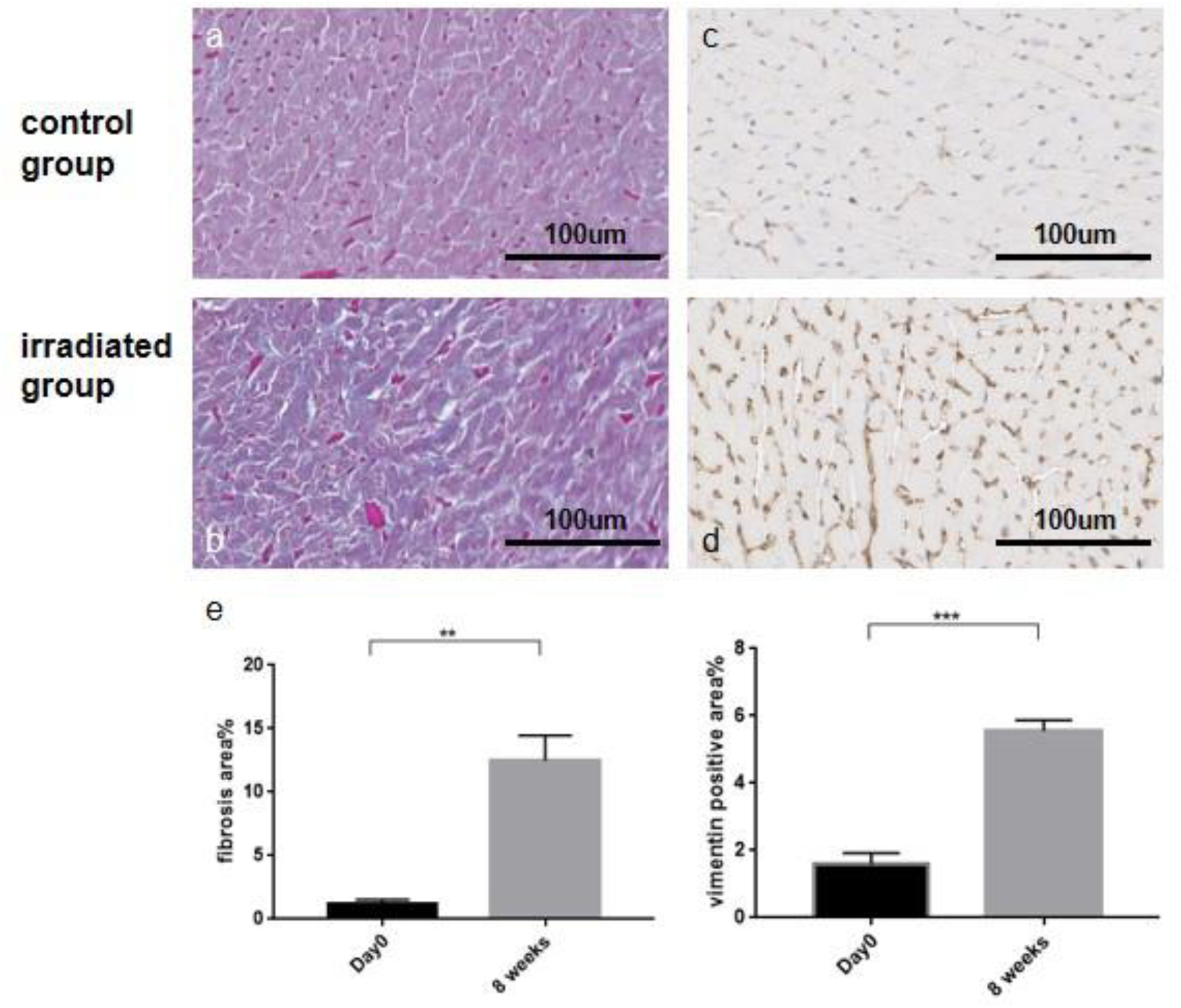
(a,b,c,d)were pathological analysis for cardiac tissue samples. (a) Masson trichrome staining for control group,(b) for irradiated cardiac tissue 8 weeks after irradiation.(c) Immunohistochemical staining for vimentin in non-irradiated sample,(d) vimentin for 8 weeks after cardiac irradiation. Irradiated tissue had more collagen deposits in the myocardial interstitium compared with control group. Vimentin was minimally expressed in non-irradiated cardiac samples, but was clearly upregulated 8 weeks after cardiac irradiation.(e) Quantitative analysis for fibrosis area and vimentin positive area. Columns represent the mean value of each group, bars±SEM. ***P<0.001, **P<0.01.(×400 microscope)

### Identification of immune cell subgroups in the peripheral blood and tissue of mice

Blood and tissue samples were analyzed respectively with 9 mice in each group. We first used viSNE to explicitly separate major immune populations and data were pooled to generate final 2-dimensional density plots. The major cell subsets were delineated in Figure 4. In tSNE plots, immune cells are clustered according to the expression profiles of marker genes(Figure 5a,b). We identified 6 major clusters in peripheral blood including neutrophil (CD45+CD11b+Ly6G+), CD4+ T cells (CD45+CD3+CD4+), CD8+ T cells(CD45+CD3+CD8+), B cell (CD45+CD19+CD24+), monocyte (CD45+CD11b+Ly6c+), macrophage (CD45+CD11b+F4/80+). In the tissue sample, we discovered 7 major clusters including endothelial cells(CD31+), neutrophil, CD4+ T cells, CD8+ T cells, B cell, monocyte and macrophage.

**Figure 4.**
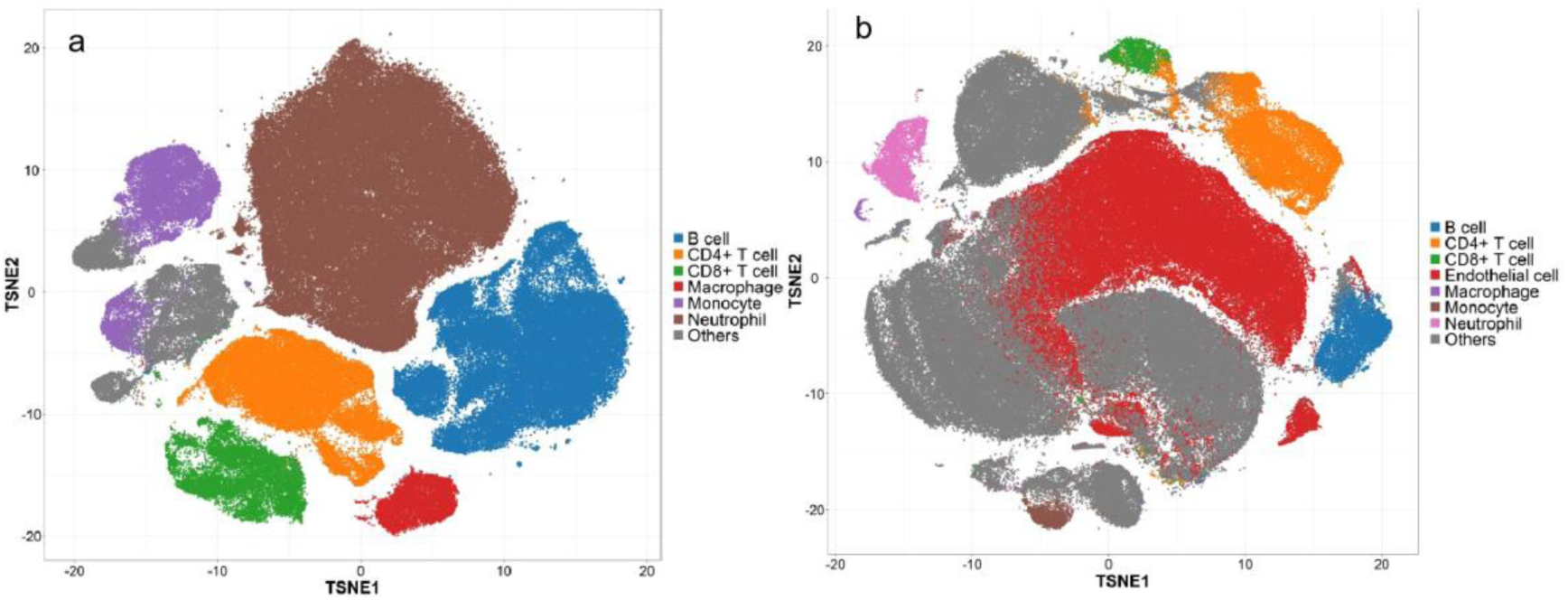
Using viSNE algorithms to identify 6 major immune cell populations.(a) Six major cell subtypes in mice blood samples, (b)Seven major cell subtypes in mice cardiac samples.

**Figure 5.**
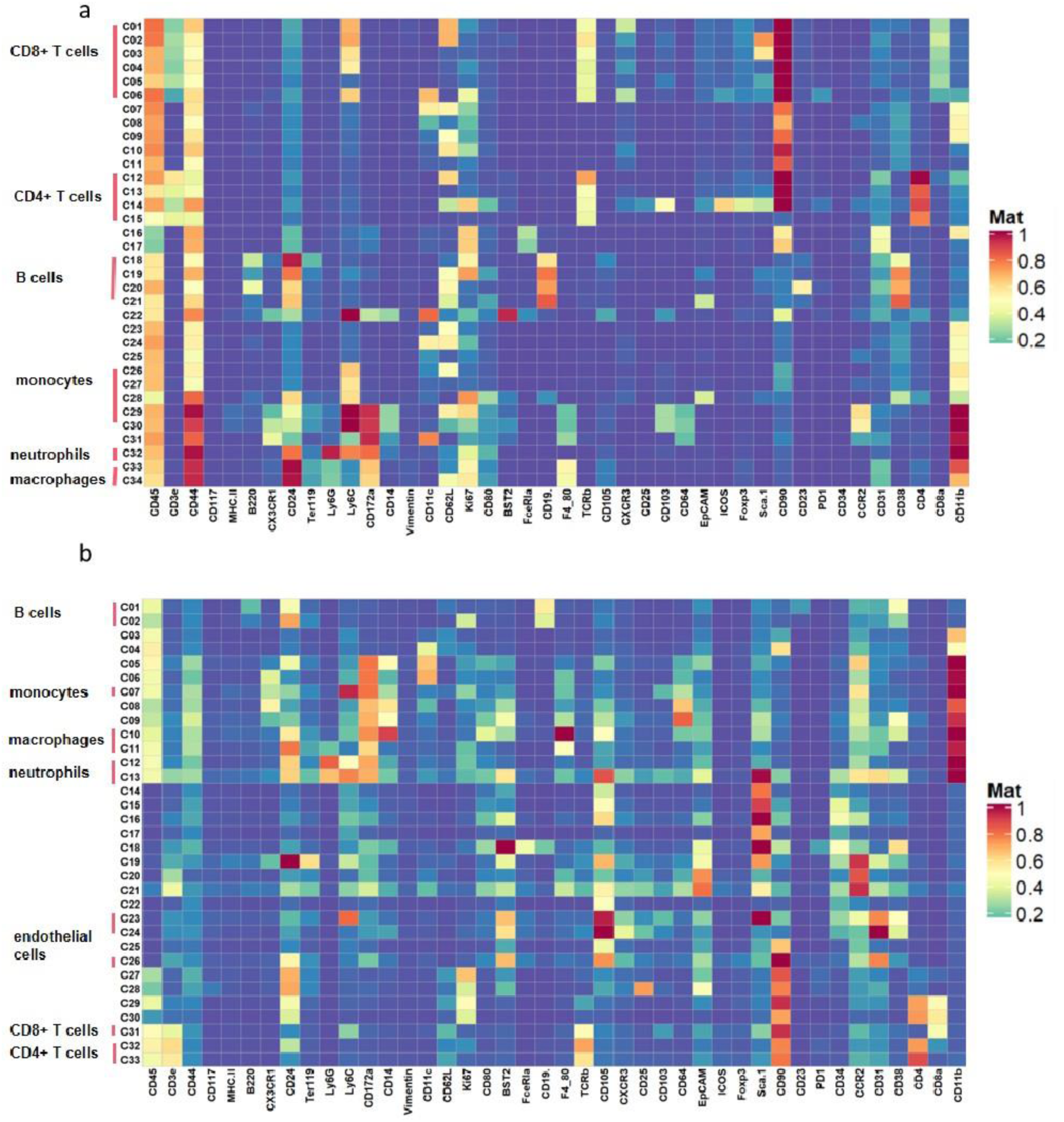
Clustering and expression level of functional markers in different cell subsets in the blood and cardiac tissue samples

### Immune cell activation in the blood after thoracic radiation

In order to identify the change of different immune cell subgroups at baseline, 7 days, 4 weeks and 8 weeks, we use bar charts to demonstrate the percentage of immune cell subgroups (Figure 6). Neutrophils accounted for the highest proportion. The results revealed that the blood samples collected after heart radiation contained a much higher proportion of neutrophils compared with samples before radiation. This increase lasted until at least 4 weeks after cardiac irradiation. The neutrophil is widely acknowledged as a key mediator in inflammatory responses and is proved to play a predominant role in the acute phase of cardiac injury following thoracic radiotherapy(11). The ratio of macrophage also increased after cardiac irradiation. Previous reports confirmed that macrophage play an important part in the process of tissue fibrosis(12). Monocyte showed similar trend as macrophage. For CD4+T cells, CD8+T and B cells, we identified significant reduction after cardiac irradiation compared with baseline samples. Past research proved that ionizing irradiation caused atrophy of the thymus, resulting in reduction of lymphocyte count and decline of lymphocyte proliferation(13). We speculated that the decline of CD4+T cells, CD8+T and B cells potentially attribute to the exposure of thymus to radiation.

**Figure 6.**
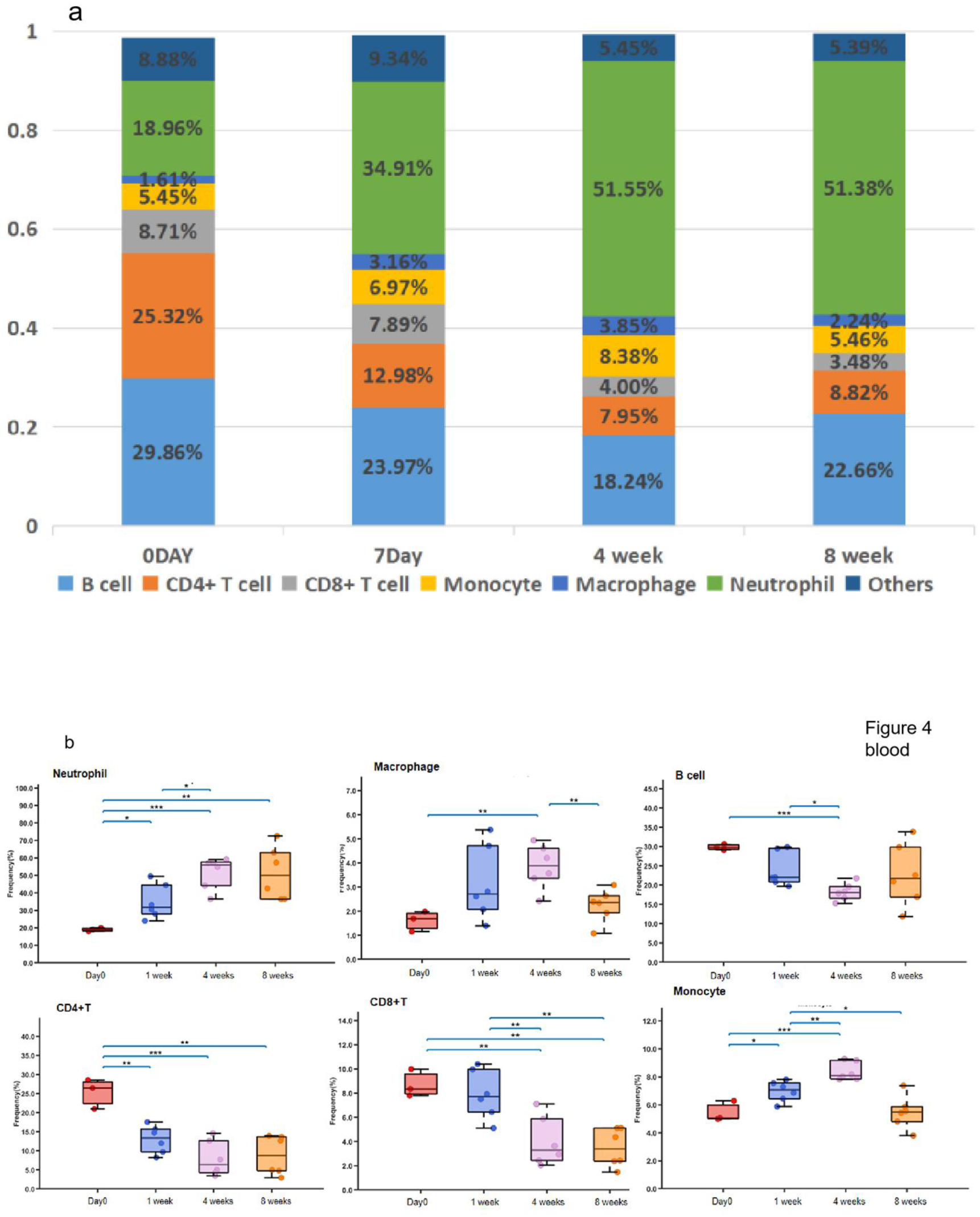
(a) Histogram of cell proportion in mice blood samples at different timepoint after cardiac irradiation.(b) Boxplot of frequencies of major subsets in the blood samples, compared by unpaired t-test. Each dot represents data from one sample. bars±SEM, *inducates p<0.05, **inducates p<0.01, ***inducates p<0.001.

### Immune cell activation in the tissue after thoracic radiation

From the Figure 7a, we discovered that endothelial cells made up the largest proportion of cells, followed by CD4+T cells. The bar chart showed that the ratio of endothelial cells in tissue samples fell considerably after heart irradiation and no sign of increase was observed later at the time of four or eight weeks(Figure 7b). Bjorn concluded that ionizing radiation can activate the caspase pathway by ceramide formation and persistent p53 signaling, causing endothelial cell death. Thus, the physiological endothelial barrier is compromised(1). The proportion of macrophage rose drastically 4 weeks after cardiac irradiation and remained high until 8 weeks. Macrophages are central players in the immune response following tissue injury. Previous investigation revealed that radiation-induced endothelial-to-mesenchymal transition (EndMT) leads to M2 polarization of macrophage(14). M2 macrophages are considered to be pro-fibrotic mainly because they are present in scars and fibrotic tissue and express pro-fibrotic factors like TGF-β1(12). The alteration of monocyte was similar as macrophage. The proportion of CD4+T and CD8+T cells within cardiac tissue elevated after 4 weeks. Past evidence supports that CD4+T cells as the primary cell type involved in the process of fibrosis through direct adhesion and actions on cardiac fibroblasts. Peripheral T cells become activated, which enables their recruitment to the left ventricle through cell surface interaction with ICAM-1 and myocardial CXCR3 ligands(13). The ratio of B cell fell 1 week after irradiation and rose after 4 weeks. As for neutrophils, the ratio at 4 weeks after irradiation tripled that at Day0, the result is similar to those previously reported for radiation lung injury(15).

**Figure 7.**
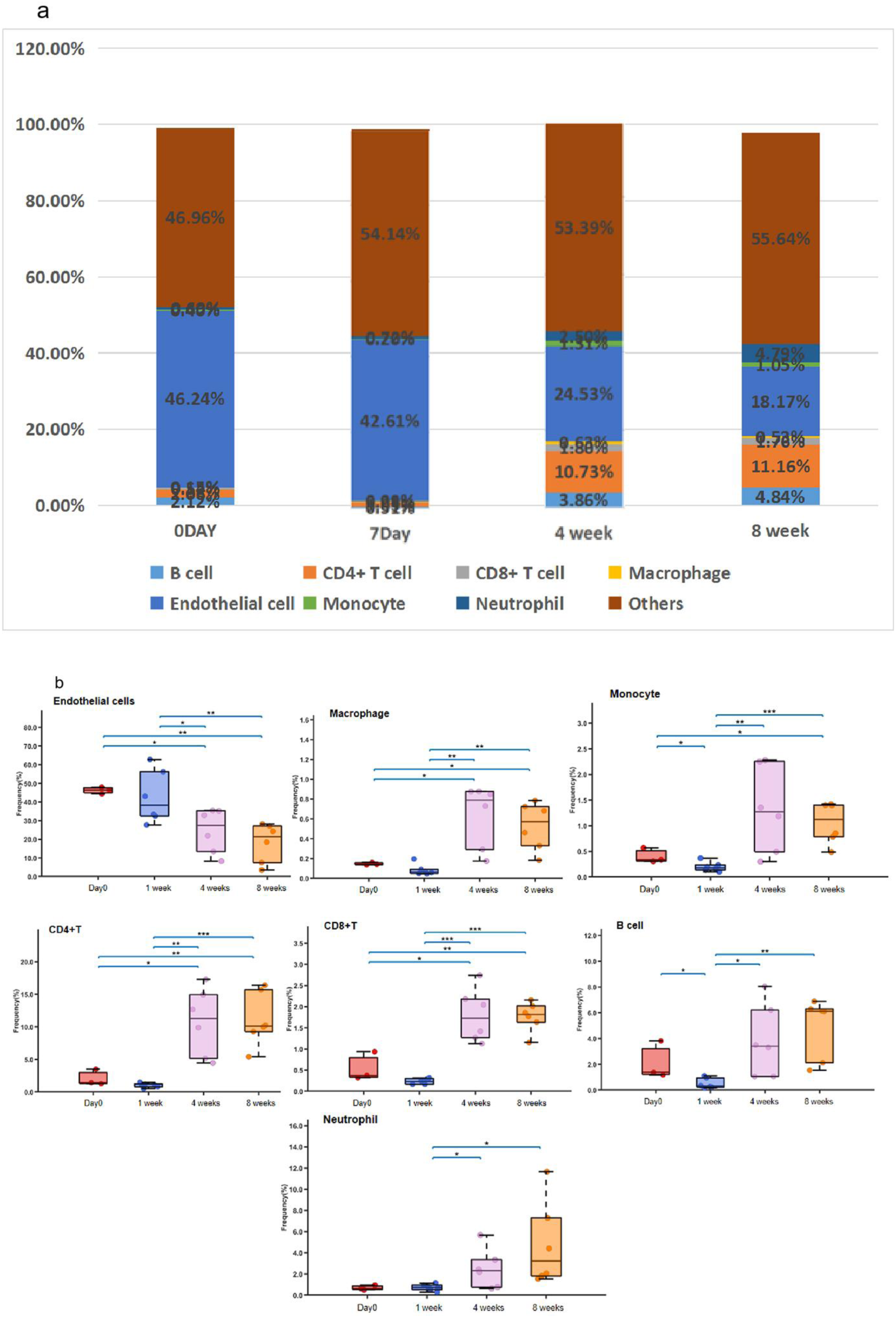
(a) Histogram of cell proportion in mice cardiac samples at different timepoint after cardiac irradiation.(b) Boxplot of frequencies of major subsets in the cardiac tissue samples. Each dot represents data from one sample. bars±SEM, *inducates p<0.05, **inducates p<0.01, ***inducates p<0.001.

### Cytokine alteration after cardiac irradiation and The role of macrophage in radiation-induced cardiac fibrosis

Cytokine expression may reflect the status of cardiac fibrosis. We determined the level of TGF-β, TNF-α, IL-6 in the serum of mice before and 4 weeks after cardiac irradiation through ELISA. All above cytokines participated in the activation of M1 macrophage and stimulation of M2 macrophage thus contribute to cardiac fibrosis(12). The results revealed that the expression levels of TGF-βat baseline 1.74±0.42 pg/ml vs. 53.69±25.50pg/ml after irradiation(p=0.0071), TNF-α 1332±62.49pg/ml at baseline vs. 1797±146pg/ml after irradiation(p=0.0064), IL-6 0.52±0.28pg/ml at baseline vs. 1.58±0.20pg/ml after irradiation(p=0.0243) were significantly enhanced(Figure 8a).

**Figure 8.**
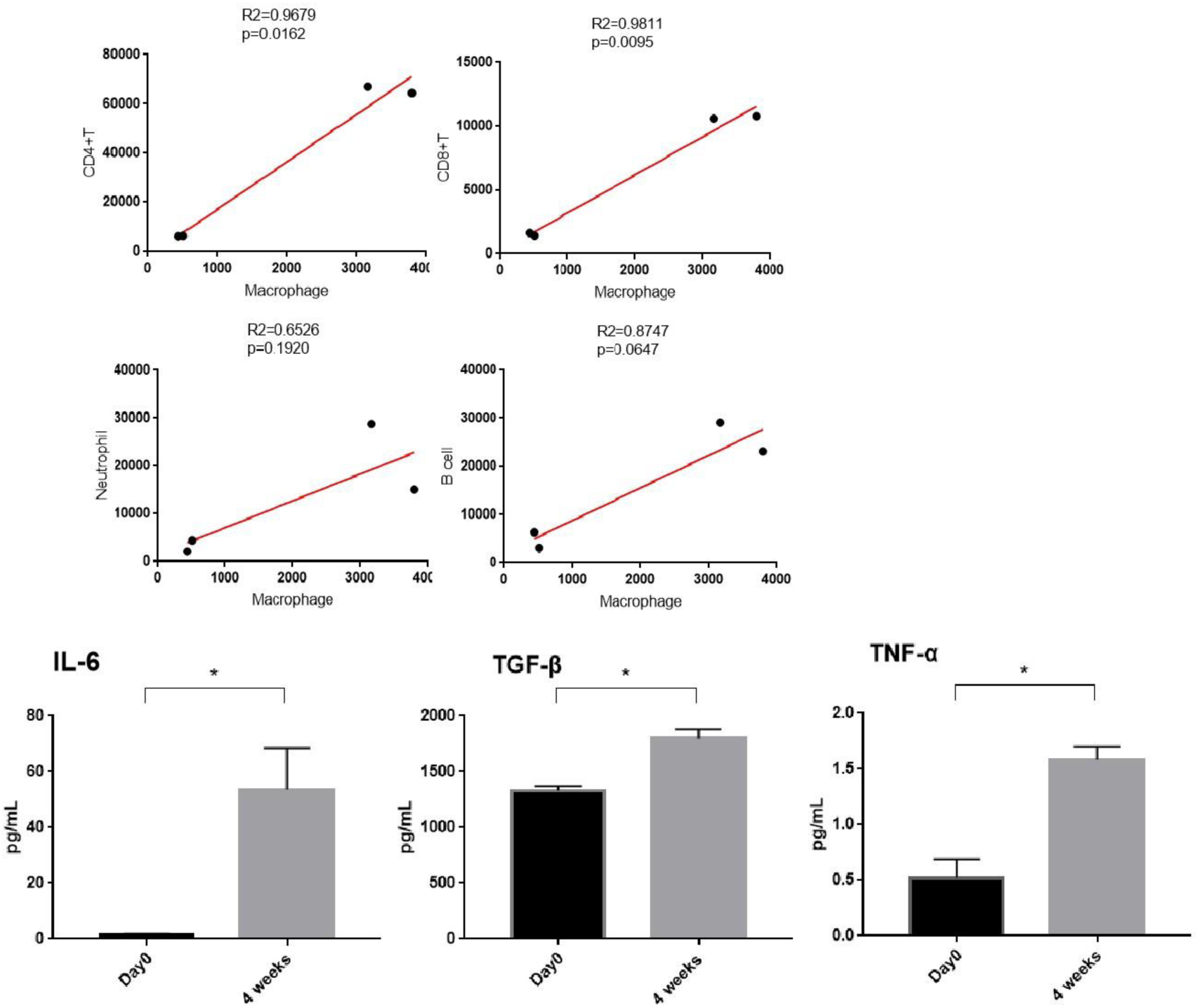
(a) The level of IL-6, TGF-βand TNF-α after cardiac irradiation compared with control group. bars±SEM, *inducates p<0.05. (b) Correlation plot of CD4+T, CD8+T, neutrophils and B cells with T cells.

Macrophage were crucial for the activation of fibroblasts, which transform to a myofibroblast phenotype and contribute to cardiac fibrosis after cardiac injury. Based on patterns of gene and protein expression and function, macrophages are commonly classified as classically activated (M1) or alternatively activated (M2) cells. M1 macrophages release pro-inflammatory cytokines, such as IL-1β, IL-6, and TNFα. M2 macrophages release anti-inflammatory factors, such as IL-10, TGF-β1 and promote immunosuppression and scar resolution(12). As TGF-β, TNF-α, IL-6 all increased after cardiac irradiation and their close relationship with the evolution of macrophage function during tissue repair, we assumed that macrophage also play a critical part in radiation-induced cardiac fibrosis. In order to explore the role of macrophage during radiation-induced cardiac fibrosis, we performed correlation plot between macrophage and other immune cells in cardiac tissue. The result demonstrated that CD4+T and CD8+T were strongly positively correlated with macrophage. Neutrophils and B cell did not show significant correlation with macrophage(Figure 8b).

## Discussion

A large portion of patients with lung cancer, esophageal cancer and lymphoma require thoracic radiotherapy. However, with the prolongation of the survival of cancer patients, more of them start to develop cardiovascular symptoms, which raises the concern of radiation-induced cardiac injury. A retrospective analysis included 748 consecutive locally advanced NSCLC patients treated with thoracic radiotherapy. After a median follow-up of 20.4 months, 77 patients developed ≥1 major adverse cardiac events(MACE), the 2-year cumulative incidence is 5.8%. Mean heart dose (≥10 Gy vs <10 Gy) was associated with a significantly increased risk of all-cause mortality (ACM) in CHD-negative patients(16). Late cardiovascular effects manifest decades after treatment and result in a variety of cardiovascular complications including the following: valvular heart disease, vasculopathy including coronary artery disease, pericardial disease, conduction system dysfunction and myocardial fibrosis(17). Although there exist substantial evidence about potential mechanism of radiation-induced cardiac fibrosis(18, 11), the interconnection of various immune cells during this process is not entirely understood.

In our study, the surviving mice undergone cardiac irradiation with SARRP platform present signs of myocardial fibrosis through pathological and functional changes. Shisuo Du et al also deliver cardiac irradiation with SARRP. Reduced cardiac EF,FS,LVID and more collagen deposits were observed in the group of irradiated heart compared with control mice, which is similar with our own findings(19), indicating we established a preliminary model of RICF in mice.

The key contribution of our study is that we identified the alteration of different cell populations in the peripheral blood and cardiac tissue samples during RICF. In order to illustrate the response towards thoracic irradiation, the cell populations were analyzed at day0 (before irradiation), 1 week, 4 weeks and 8 weeks (after irradiation). We found that the proportion of neutrophils increased considerably after thoracic radiation in the peripheral blood and tissue samples. After cardiac irradiation, damage to vascular endothelium stimulates the release of chemokines, which permeabilise the vessel wall and promote recruitment and proliferation of inflammatory cells(2). Neutrophils are associated with acute myocardial inflammatory responses and contribute to tissue repair in the early phase of radiation-induced cardiac fibrosis(20). Sohn et al explored that mice exhibited higher numbers of neutrophils, lymphocytes, eosinophils, macrophages compared to controls after thoracic irradiation in the bronchoalveolar lavage fluid(21). Contreras et al discovered increased rates of immunosuppression and elevated level of neutrophil-to-lymphocyte ratio(NLR) after definitive treatment of locally advanced non-small-cell lung cancer (LA-NSCLC) with radiotherapy(22). Collectively, these results suggest that cardiac irradiation can contribute to the elevation of neutrophil and trigger acute inflammatory response.

We also discovered that the proportion of macrophages rose greatly after cardiac irradiation compared with control groups. Macrophage is one of the major components of the accumulated inflammatory infiltration after irradiation in tissue. Early in response to tissue damage, neutrophils migrate to the injured area, and macrophages then engulf these apoptotic neutrophils. Depending on the surrounding microenvironment, macrophages are phenotypically able to switch to an anti-inflammatory M2 phenotype(12). M2 macrophages have often been considered to be pro-fibrotic because they present in scars and fibrotic tissue and express profibrotic factors, particularly TGF-β1. Moreover, macrophages facilitate the proliferation and activation of myofibroblasts, which are the primary effectors of the heart’s fibrotic response(23). Also, macrophages release matrix metalloproteinase(MMP), which is an important enzyme that degrades extracellular matrix and accelerate the process of fibrosis(24).

As for cytokine measurement, we found that IL-6, TGF-βand TNF-αincreased considerably in the peripheral blood after cardiac irradiation. Endogenous TGF-βplays an important role in the pathogenesis of cardiac fibrotic and hypertrophic remodeling(25). TGF-βstimulation induces myofibroblast differentiation and enhances extracellular matrix protein synthesis. In addition, TGF-β exerts potent matrix-preserving actions by suppressing the activity of MMP and by inducing synthesis of protease inhibitors(25). IL-6 possesses a wide range of biological activities in immune regulation, hematopoiesis, and inflammation. Giselleet et al. reported that IL-6 infusion cause myocardial fibrosis, hypertrophy, and diastolic dysfunction in rats(26). Past research identified that inhibiting IL-6 gene expression can prominently ameliorate myocardial infarction(MI)-induced myocardial remodeling(27). Evidence suggests that TNF-α promotes cardiac fibrosis, triggering a predominant matrix-degrading phenotype. Fibrotic remodeling of the TNF-αoverexpressing heart is correlated with increased expression of TGF-βand has been suggested to involve interactions between fibroblasts and mast cells(20). In conclusion, IL-6, TGF-βand TNF-α are secreted in the cardiac interstitium and play distinct roles in activating specific aspects of the fibrotic response.

Macrophages are capable of producing and secreting large amounts of pro-inflammatory mediators such as the interleukin IL-1β, TNF-α, IL-6, and pro-fibrotic growth factors such as TGF-β, platelet-derived growth factors(PDGFs), and fibroblast growth factors(FGFs)(41). Thus we speculate the vital role of macrophage in the course of radiation-induced cardiac injury. Previous results verified that the resolution of scarring and fibrosis appears to be the responsibility of macrophages(28,29,30). At early stages of external stimulus, macrophages respond to the presence of PAMPs, DAMPs, and/or Th1 effectors and become classically activated (M1) cells, which are characterized by pro-inflammatory mediators and potent anti-microbial activity. Next, with the effect of Th2 mediators such as IL-10, IL-13, the alternative activation (M2) of macrophages were activated and possess anti-inflammatory, pro-angiogenic and pro-fibrotic properties(12, 31).

As macrophages play critical roles during the initiation, maintenance, and resolution phases of tissue repair, we tried to explore the interconnection between macrophages and other immune cells. We discovered that the population of CD4+T and CD8+T were positively correlated with macrophages according to the correlation plot. TA Wynn concluded many experimental models of fibrosis and discovered that CD4+ T cells play a prominent role in the process(32). T-cell infiltration of the myocardium is necessary to produce LV fibrosis and dysfunction indicating a functional role for T-cells in cardiac fibrosis(4). T-cells can act through direct adhesion and actions on cardiac fibroblasts. Besides, T cells interact with macrophages, fibroblasts, and other tissue resident cells to accelerate inflammation and to promote fibrosis under certain circumstances. Different T-cell subsets can secrete several mediators and growth factors that influence the myocardial molecular milieu and directly interfere with the macrophages’ and fibroblasts’ activity(33). For neutrophils and B cells in our study, we did not find significant correlation with macrophage. However, Epelman et al concluded that after cardiac injury, when increased levels of apoptotic tissue neutrophils are sensed, macrophages reduce their own secretion of IL-23, which decreases IL-17a levels, and thereby decrease neutrophil production(34). In sum, macrophage subpopulations are sources of inflammatory and fibrogenic mediators with distinct properties that mediate pro-inflammatory, anti-inflammatory, or fibrogenic actions. More work needs to be done on the role of fibrogenic macrophage subsets in the fibrotic myocardium.

However, our study has some limitations. First, as a late-stage abnormality for radiation-induced cardiac injury, eight weeks for cardiac fibrosis may not be long enough to observe pathological and functional changes. The reason of death remains unclear. The possible reasons of death include undefined cardiac injury and lung injury caused by thoracic irradiation. Creating a model with decreased lung dose would be highly recommended. Moreover, as we did not include relevant surface markers in our panel, different macrophage groups were not able to be identified.

## Conclusion

In summary, the present study established a preclinical murine radiation-induced cardiac injury model and use mass cytometry to illustrate the alterations of cell population before and after cardiac irradiation in the blood and tissue samples. The level of several pro-inflammatory and pro-fibrotic cytokines increased considerably after irradiation. As those cytokines are all closely correlated with the evolution of macrophage function during repair, we conceived that macrophage may play a critical role in the radiation-induced cardiac injury. We also explored that the number of CD4+ and CD8+T cells were positively correlated with macrophages. Such insight should prompt the development of macrophage-targeted therapies for radiation-induced cardiac fibrosis. More detailed work needs to be done on the mechanism of RICF in order to reduce cardiac complications after thoracic radiotherapy.

## Author contributions

I. Conception and design: Bing Xia, Mingliang You, Xue Zhang
II. Administrative support: Bing Xia, Mingliang You
III. Provision of study materials or patients: Bing Xia, Yi Tang
IV. Collection and assembly of data: All authors
V. Data analysis and interpretation: All authors
VI. Manuscript writing: All authors
VII. Final approval of manuscript: All authors

## Acknowledgments

We appreciate Zhejiang Chinese Medical University Laboratory Animal Research Center for providing animal housing and husbandry. We also thank small-animal radiation research platform (SARRP) in the Zhejiang Key Laboratory of Radiation Oncology for mice cardiac irradiation.

## Footnote

***Ethical statement:*** The authors are accountable for all aspects of the work in ensuring that questions related to the accuracy or integrity of any part of the work are appropriately investigated and resolved. All animal studies were performed in accordance with an animal use protocol approved by the Animal Care ethics board of Zhejiang Chinese Medical University.

---

**Supplementary Table 1.** 42 cell lineage marker to identify different cell populations

| Number of marker | Marker | Number of marker | Marker |
| --- | --- | --- | --- |
| 1 | CD45 | 22 | F4/80 |
| 2 | CD3e | 23 | TCRβ |
| 3 | CD44-m/h | 24 | CD105 |
| 4 | CD117 | 25 | mCD183(CXCR3) |
| 5 | MHC II | 26 | CD25 |
| 6 | B220(CD45R) | 27 | CD103 |
| 7 | CX3CR1 | 28 | CD64 |
| 8 | CD24 | 29 | EpCAM |
| 9 | Ter119 | 30 | ICOS |
| 10 | Ly6G | 31 | Foxp3 |
| 11 | Ly6C | 32 | Sca-1 |
| 12 | CD172a(SIRPα) | 33 | CD90(Thy-1) |
| 13 | CD14 | 34 | CD23 |
| 14 | Vimentin | 35 | PD1,cd279 |
| 15 | CD11c | 36 | CD34 |
| 16 | CD62L | 37 | CCR2 |
| 17 | Ki67 | 38 | CD31 |
| 18 | CD80 | 39 | cd38 |
| 19 | BST2 | 40 | CD4 |
| 20 | FcεRIα | 41 | CD8a |
| 21 | CD19 | 42 | CD11b |

